# Ultrasound mediated cellular deflection results in cellular depolarization

**DOI:** 10.1101/2021.06.11.447976

**Authors:** Aditya Vasan, Jeremy Orosco, Uri Magaram, Marc Duque, Connor Weiss, Yusuf Tufail, Sreekanth H Chalasani, James Friend

**Affiliations:** Medically Advanced Devices Laboratory, Department of Mechanical and Aerospace Engineering, Jacobs School of Engineering and Department of Surgery, School of Medicine, University of California San Diego, La Jolla CA 92093 USA; Molecular Neurobiology Laboratory, The Salk Institute for Biological Studies, La Jolla, CA 92037

**Keywords:** ultrasound, neuromodulation, digital holographic microscopy, acoustofluidics

## Abstract

Ultrasound has been used to manipulate cells in both humans and animal models. While intramembrane cavitation and lipid clustering have been suggested as likely mechanisms, they lack experimental evidence. Here we use high-speed digital holographic microscopy (to 100-kHz order) to visualize the cellular membrane dynamics. We show that neuronal and fibroblast membranes deflect about 150 nm upon ultrasound stimulation. Next, we develop a biomechanical model that predicts changes in membrane voltage after ultrasound exposure. Finally, we validate our model predictions using whole-cell patch clamp electrophysiology on primary neurons. Collectively, we show that ultrasound stimulation directly defects the neuronal membrane leading to a change in membrane voltage and subsequent depolarization. Our model is consistent with existing data and provides a mechanism for both ultrasound-evoked neurostimulation and sonogenetic control.

## 1 Introduction

Existing methods to stimulate neural activity include electrical [1, 2, 3, 4, 5], optical [6] and chemical techniques [7]. They have enabled the development of novel therapies that are used in clinical settings [8], in addition to helping understand aspects of neural function [9] and disease mechanisms [10]. Despite their beneficial impact, these approaches are fundamentally limited. Electrical stimulation is invasive, requiring direct contact with the target of interest. Inserting electrodes into the brain may lead to inflammation, bleeding, cell death [11], and local cytokine concentration increases in microglia that precipitate astrocyte formation around the electrodes that, in turn, reduce long-term effectiveness [12]. In addition, it may have non-specific effects depending on the electric field generated by the electrodes and the stimulation parameters used [13]. Transcranial direct current stimulation (tDCS) and transcranial magnetic stimulation (tMS) are new and non-invasive, yet they have poor spatial resolution on the order of 1 cm [14, 15]. Furthermore, approaches combining genetic tools with light or small molecules achieve cellular specificity. *Optogenetics*, which involves the use of light and genetically encoded membrane proteins [16], has enabled elucidation of cellular circuits in animal models. However, it remains an invasive technique and applications are limited by the depth of penetration of light in tissue. In contrast, *chemogenetics*, using small molecule sensitive designer receptors, is limited by poor temporal resolution and is unfortunately impractical for many neural applications that require millisecond response times [17].

Ultrasound can overcome the limitations of these methods. It is non-invasive and has a high spatiotemporal resolution. Improvements in the spatial resolution through transfection currently come at the cost of a minimally-invasive procedure to directly inject the vector into the target tissue [18], though there may soon be non-invasive alternatives [19]. The spatial resolution of ultrasound is governed by the wavelength of operation and is about 1.5 mm at 1 MHz in tissue. The temporal resolution is dependent on the duration of stimulation and may be as short as a single time period, *T=*1/ *f* where *f* is the operating frequency. The frequency is dictated by the depth and size of the target region in traditional focused ultrasound neuromodulation [20], and is generally compatible with human and animal anatomy for this application. Harvey [21] was one of the first to utilize these advantages over ninety years ago on frog ventricular heart tissue. Recent advances in describing the suppression of epileptic activity in patients [22] are an indicator the method is still being considered in clinical applications.

Despite these recent experimental and clinical developments, and progress in exploring the sonogenetic and ultrasonic-to-chemical action mechanisms, there is no convincing, overarching explanation for the observations. Some of the proposed mechanisms include cavitation [23], indirect auditory signalling *in vivo* [24] and increased lipid clustering resulting in a change in the membrane tension [25]. These studies have either been conducted on time scales that are orders of magnitude larger than those used for ultrasound neuromodulation or lack robust imaging techniques that operate at timescales relevant to the frequency of stimulation.

Additionally, studies often treat surface tension, membrane composition, and membrane stresses as a single term, *membrane fluidity* [25]. This term lacks rigourous physical description and is assigned a value based on relative fluorescence intensity changes. The imprecision of this description makes it difficult to isolate the influence of the measurable physical mechanisms of which it is comprised. A model using membrane fluidity leaves the explanation of the biophysical phenomenon incomplete.

More broadly, action potentials are known to accompany, in phase, the cell membrane’s deflection [26, 27]. These observations and more recent studies into the thermodynamic effects associated with the generation of action potentials [28] point to transmembrane voltage changes being more than just an electrical phenomenon. In addition, phenomenological observations of ultrasound on neurons suggests there is a conversion of mechanical to electrical energy at the level of individual neurons.

The model described here details membrane deflection due to an applied ultrasound stimulus and links it to neuronal depolarization. In addition, we report the first three-dimensional visualization of cell membrane deflection due to an ultrasound stimulus by using high-speed digital holographic microscopy (DHM). We use current clamp electrophysiology in the challenging environment of intense ultrasound to monitor ultrasound-driven, real-time changes in voltage across the membrane in single neurons *in vitro*. Our experiments confirm the predictions made by the biophysical model, both with regard to membrane deflection and voltage changes. Our findings provide insight into the effects of ultrasound on cells and cell signaling, the understanding of which is vital to sonogenetics and its clinical application.

## 2 Results

### 2.1 Digital holographic imaging of cell membrane deflection

High resolution imaging approaches employing phase-contrast [29] and differential contrast [30] are commonly used to image biological specimens. These techniques transform phase differences to amplitude differences in an image, but they lack quantitative phase information. High-speed digital holographic microscopy (DHM) [31] is a cutting-edge method that yields three-dimensional holograms at high frame rates. We use transmission DHM, which measures transparent media based on quantifying phase disparities induced by the measured sample. In short, this approach works by comparing phase differences induced in the coherent light transmitted through the sample with reference light traversing an unobstructed path. Digital holographic microscopy has several advantages in comparison to conventional microscopic techniques. Numerical processing of the wavefront transmitted through the sample permits simultaneous computation of intensity and phase distribution [32]. The holographic measurements also make it possible to focus on different object planes without relative movement between the stage and the lens [33] and enable numerical lens aberration correction [34].

The response measurements consisted of a 25 ms baseline recording, followed by a 50 ms stimulus and a 25 ms post-stimulus dwell, leading to a median deflection of 214 nm for human embyronic kidney (HEK293) cells and 159 nm for neurons, with a range of 100 nm to 550 nm across the two tested cell types (Supplementary Videos 1 and 2). The baseline deflection had a range of ±20 nm, inclusive of both random thermal fluctuations across the cell membrane and noise introduced to the system due to the imaging arrangement (Fig. 1b). The deflection predicted by the model depends on the length of the membrane and surface tension. The values predicted in Fig. 2b are representative of typical cell sizes used in our experiments for HEK cells and neurons. Sample displacement profiles and membrane profiles are illustrated in Fig. 1c-d (*see* Supplemental Videos as well), and Fig. 1e represents the deflection profile for a cluster of neurons. The cluster was imaged to confirm deflection in a group of neurons and help provide insight into the *in vivo* mechanisms of activation.

**Figure 1:**
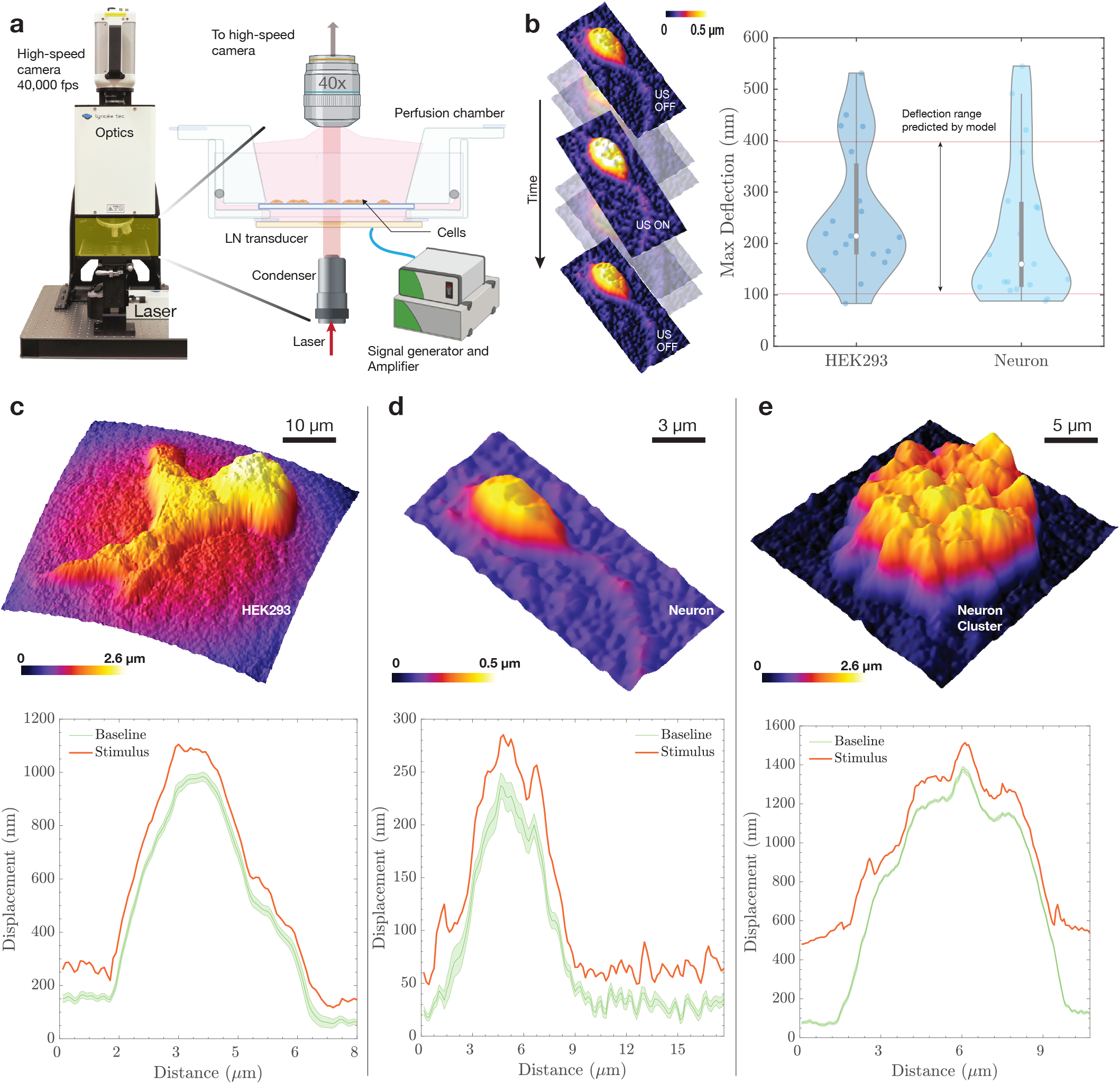
High-speed DHM imaging of membrane deflection. The deflection predicted by the model was verified using **(a)** high-speed digital holographic microscopy. The DHM setup consists of a lithium niobate transducer operating at 6.72 MHz driven by a signal generator and an amplifier. The cells are mounted on a coverslip and placed in a custom perfusion chamber maintained at 37°C. The DHM enables the **(b)** quantitative reconstruction of phase images acquired by the high-speed camera recording at up to 200,000 frames per second. The maximum deflection is confirmed to be within the range predicted by the model (100–400 nm), with a median deflection of 214 nm for HEK293 cells and 160 nm for neurons. All recordings invovle a 25 ms baseline, followed by a 50 ms burst and a 25 ms post-stimulus dwell. The displacement profiles of **(c-e)** different cell types studied indicate a steady baseline reading, with a maximum variation of ±20 nm and the maximum deflection similar across the studied cell types.

**Figure 2:**
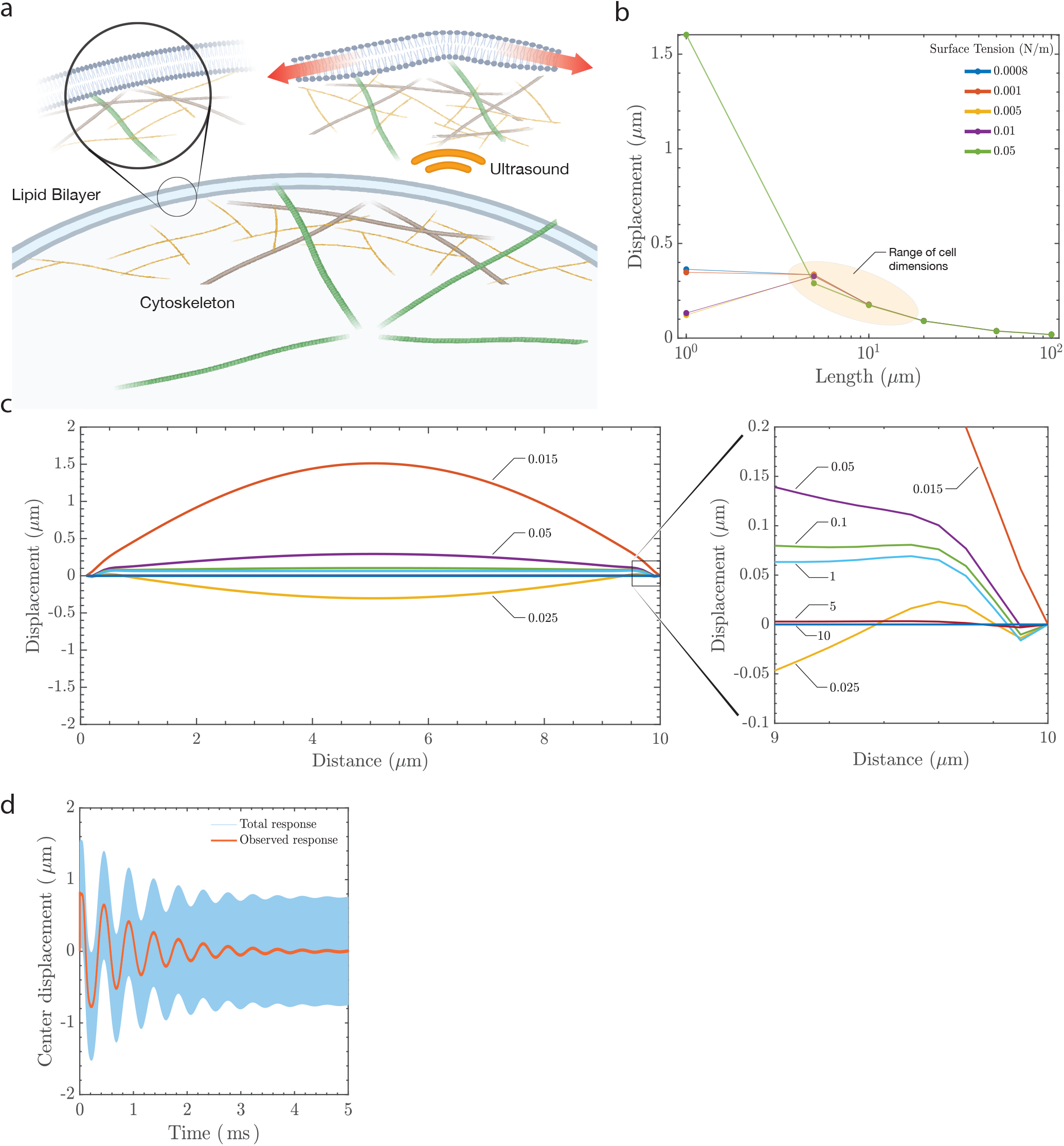
Prediction of membrane deflection due to ultrasound. Ultrasound results in (**a)** membrane deflection that triggers a transmembrane electrical response. The cell membrane bilayer stretches, increasing its area, and the outer leaflet of the bilayer will likely deflect more than the inner leaflet due to the the presence of cytoskeletal components such as actin and microtubules that anchor the inner leaflet. Two of the factors that affect membrane displacement are surface tension of the lipid membrane and the length under consideration. The model **(b)** predicts displacements between 100–400 nm for dimensions that correspond to the size of a cell (5–20 µm). The response is **(c)** dynamic, with snapshots of the predicted deflection at different times (in *ms*) across a 10 µm wide membrane section that is anchored at the ends. The maximum deflection occurs when the stimulus is first provided. A low-pass temporal filter of the membrane’s center displacement at 5 µm indicates **(d)** an oscillatory deflection over at least 5 ms.

The length of the membrane under consideration can vary depending on the cell type and distance between focal adhesions, and, as later predicted by the model, the length of the membrane between the fixed points and the surface tension under consideration are crucial in determining the deflection (Fig. 2b). Results from the neuronal cluster show that the magnitude of deflection remains roughly the same for a group of cells as for a single neuron.

Membrane deflection during the generation of action potentials has been observed in the past [35], but the converse phenomenon of membrane deflection leading to the generation of action potentials has not been explored. Other imaging techniques have been reported for measuring cell membrane deflection, such as atomic force microscopy (AFM). Although AFM is well established and offers high spatial resolution, it suffers from low temporal resolution and lacks the ability to simultaneously scan multiple points [36]. Overall, our experimental setup allows us to confirm membrane deflection due to ultrasound and the results are within the range of membrane deflections predicted by our model.

### 2.2 Membrane deflection model

Based upon the results from the experiments, with cells cultured on a surface and surrounded by media, the membrane is assumed to be fixed at the periphery. A similar case occurs *in vivo*, where the extracellular matrix holds individual cells in place and provides anchoring locations for sections of the membrane. Cellular anchoring is important because it imposes a characteristic distance over which the range of permissible deflection wavemodes may occur [see methods]. Its deflection is restricted in the analysis to a single direction, perpendicular to the plane of the membrane and parallel to the direction of propagation of sound. The model does not take into account the restoring effects of the actin cytoskeleton, difficult to estimate but likely playing a crucial role in restoring the membrane to its original equilibrium position.

The stimulus provided to the cells is in the form of a *burst*, a short-term continuous ultrasound signal of constant amplitude and frequency. In a burst, a sinusoidal electrical signal is typically applied across the piezoelectric material used in a transducer, which transforms this signal into a sinusoidally varying pressure field in the fluid medium at the frequency of excitation. This is rather different than the approach used by Prieto et. al [37], where the ultrasound is modeled as a step increase in hydrostatic pressure from zero to a fixed positive value at *t=*0. In our approach [see methods], the burst signal oscillates at the ultrasound frequency, and an analytical solution for the slower time scale of the membrane mechanics is found in response to this harmonic ultrasound excitation. This solution is then used in a numerical model to produce the solution for the deflection of the fixed membrane, resolving the discrepancy between the timescales of ultrasonic stimulation (∼0.1 *µ*s) and the experimentally verified membrane deflection occurring on the order of milliseconds. This hybrid approach was chosen because a numerical simulation of the entire phenomena from ultrasound to membrane deflection would be extremely difficult due to the vastly different spatiotemporal scales, even with state-of-the-art computational resources. Finally, the hydrostatic pressure included by Prieto et. al [37] is discarded here, because it is orders of magnitude lower than the ultrasonic radiation pressure.

The damped wave equation describing the deflection, *u*, of the membrane in response to ultrasonic pressure, *P*_US_, is written as

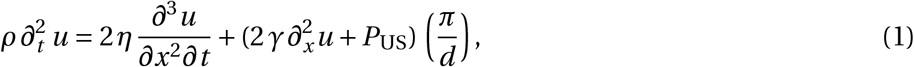

where *ρ* and *η* are the dynamic viscosity and density of the surrounding fluid, both assumed to be the same as water as used in prior studies [38, 39]); *γ* is the surface tension between the membrane and media; and *d* is the characteristic length of the membrane between anchor points. Equation (1) was solved by the method of eigenfunction expansion [methods]. Figure 2 provides results representative of the analysis, with a 1 MPa pressure supplied to the membrane using a 7 MHz transducer in the form of a sine wave over a period of 5 ms. The mechanical index for the parameters listed in this study is 0.37, well below the oft-cited mechanical index threshold for cavitation onset of 0.7 in bubble-perfused tissue [40]. However, our study uses no bubbles and the mechanical index of 0.37 is much lower than the U.S. Federal Drug Administration’s mandated clinical safety threshold index of 1.9 without introduced microbubbles [41, 42] is more appropriate. These data suggest that we are unlikely to cause cavitation.

Maximum membrane deflection occurs when the ultrasound stimulus is applied, followed by decay due to viscous losses to the host medium. The magnitude of deflection depends on the stimulation frequency and peak pressure, with lower frequencies and higher pressures producing greater membrane deflection. The critical parameters that influence the deflection magnitude are the characteristic membrane anchor length and surface tension, as shown in Fig. 2b. The deflection predicted by the model for dimensions relevant to the size of a cell are between 100 nm to 400 nm, irrespective of the value of surface tension. We modelled membrane deflection due to a range of surface tension values reported in the literature [37, 43]. Maximum membrane deflection occurs at the midpoint of the axisymmetric membrane model. This is portrayed in Fig. 2c, where we provide graphical “snapshots” of the ultrasonically-forced membrane over time. The closed-form displacement solution to Eq. (1) allows us to link the fast ultrasonic timescales (on *µ*s order) to phenomena occurring at observable timescales (on ms order), as shown in Fig. 2d. The character of the membrane “slow time” response—that is, its ability (or lack thereof) to sustain oscillations—is governed by the value of the Ohnesorge number, *Oh*.

The nondimensional parameter *Oh* characterizes the importance of dissipative viscous forces relative to the combined interaction of conservative inertial and surface tension forces. In other words, *Oh* characterizes, on average, the extent to which the membrane dissipates or conserves mechanical energy. Typical *Oh* values for neurons range from ∼0.06 to ∼ 0.45. This implies that inertial and surface tension forces dominate over viscous forces: the slow time membrane response is characteristically oscillatory. This behavior results from the membrane’s tendency toward retaining mechanical energy in the form of sustained oscillations when 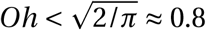. This is explicitly derived in the detailed analysis [methods] and suggests that the slow time oscillations of the ultrasonically actuated membrane is implicated in the changes in the membrane capacitance as detailed in the following sections.

### 2.3 Model prediction of action potentials and electrophysiology

Modeling the electrical output of a neuron under the influence of ultrasound involves defining a modified version of the original Hodgkin-Huxley equations [44],

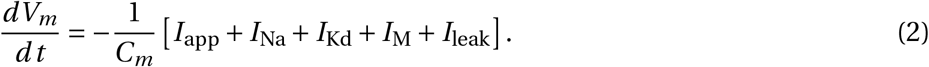

In this equation, the membrane potential of the neuron, *V*_*m*_, changes over time with respect to the membrane capacitance, *C*_*m*_, and the underlying currents, *I*_app_, *I*_Na_, *I*_Kd_, *I*_M_, and *I*_leak_. At rest, *V*_*m*_*=* −71.9 mV is the well-known membrane potential of the cell and, notably, the action potential generation is controlled by the presence of an applied current, *I*_app_, while the other currents are based on the membrane morphology and chemistry and are detailed in the methods. The increase of *I*_app_ beyond a certain threshold produces spiking behavior typical of neurons.

The capacitance, *C*_*m*_, may also fluctuate due to a morphological change in the membrane. Such a modification is not modeled in the original representation of this equation, but it may be included. The voltage change as described in eqn. (2) includes a time-dependent capacitive current, 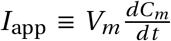. With this included in eqn. (2), it is possible to solve the differential equation for the voltage and gating variables while incorporating the capacitance change due to membrane deflection. Membrane deflection is constrained to a certain extent due to parts of the cell that are adherent to the substrate or the extracellular matrix. This causes an increase in area between the adherent locations and with sufficient deflection, this produces a depolarization across the membrane. The value of the transmembrane voltage is dependent on the magnitude and duration of the applied stimulus. Figure 3 indicates the change in capacitance due to 6.72 MHz ultrasound at 0.5 MPa (Fig. 3a) and 1 MPa (Fig. 3b) with the corresponding area fluctuations that bring about the change in capacitance represented in Fig. 3c. In order to compute the time-dependent membrane area variation, we extract the slow time output of Eq. (1) for use with the axisymmetric area integral. The capacitance of the membrane is then determined by treating it as a dielectric between charged surfaces. This yields a slow time capacitive response, bearing order of magnitude equivalence to the ion channel relaxation times in the modified Hodgkin-Huxley model [45].

**Figure 3:**
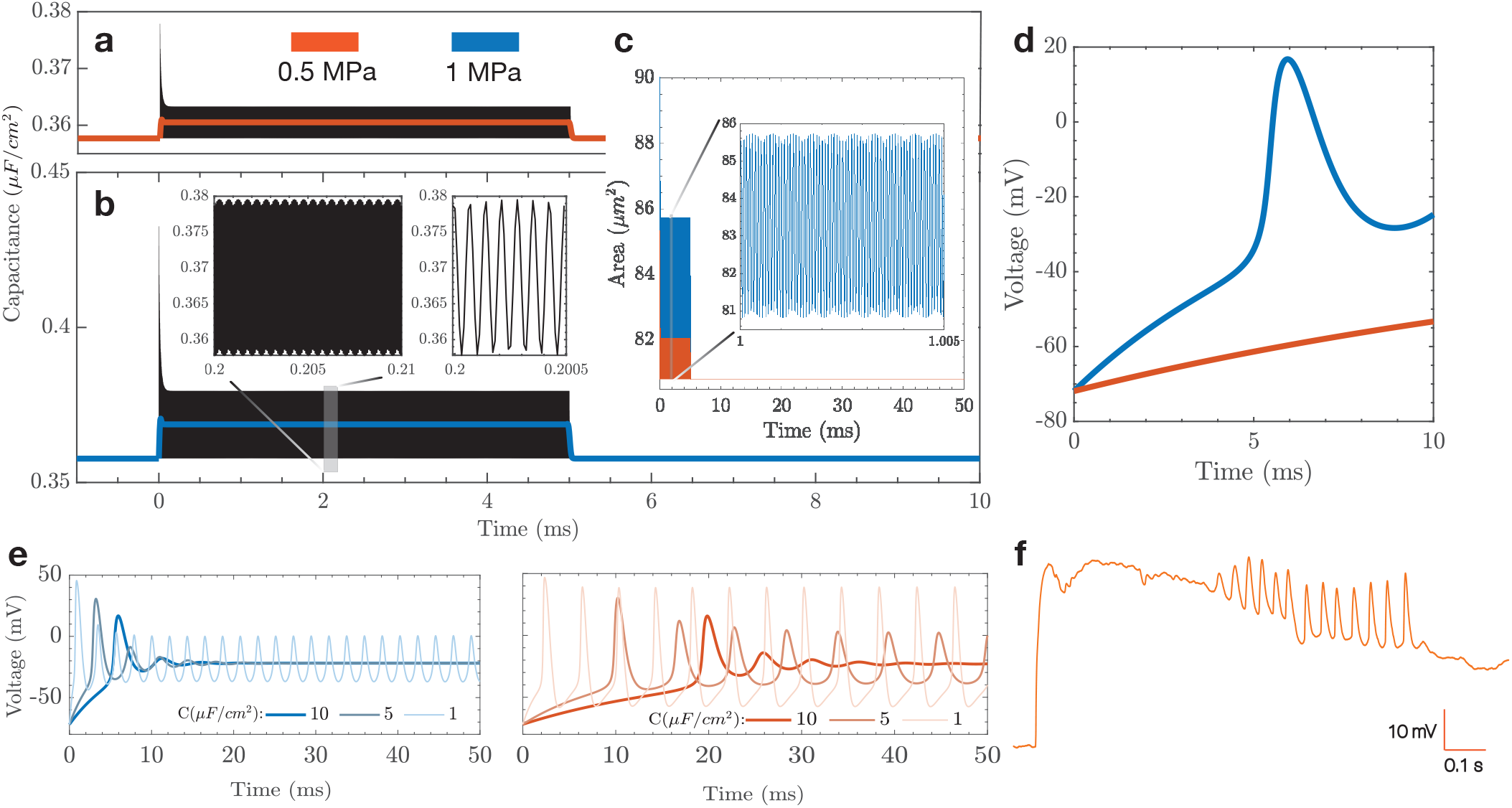
Displacement-driven capacitance changes result in action potential generation. The mechanism of ultrasound driving **(a**,**b)** capacitance changes over the stimulus duration (5 ms) for **(a)** 0.5 MPa and **(a**,**b)** 1.0 MPa. The capacitance changes occur principally due to **(c)** area changes due to membrane deformation. The capacitance change produces **(d)** depolarization in the 1 MPa case, yet fails to do soat 0.5 MPa, indicating the presence of a pressure threshold required to stimulate neurons. A longer stimulus of **(e)** 50 ms duration shows the time evolution of action potentials for the two different pressure conditions. This predicts that longer stimulus durations may be necessary at lower pressures in order to produce action potentials. Despite the challenges of retaining the clamp in the presence of ultrasound, **(f)** *in vitro* current clamp electrophysiology was used to verify the predictions of the model and shows the presence of a preliminary spike followed by oscillations in voltage across the membrane corresponding to the predictions of the model.

The stimulus of 1 MPa results in depolarization as indicated in Fig. 3d, while the lower pressure does not result in the generation of an action potential over the stimulus duration. Reported values of baseline membrane capacitance have been shown to vary [46], and we show that longer stimuli will result in the generation of action potentials as a cumulative effect of capacitance change over the duration of the stimulus. Figure 3e represents transmembrane voltage changes for a stimulus of 50 ms. We notice that depolarization takes place in both cases. However, initial spikes are delayed by up to 20 ms in the lower pressure case, indicating the need for increased stimulus durations for lower pressures. Our model also shows a lower spike frequency for the 0.5 MPa case in comparison to 1 MPa. The simulation output of our model for the lower pressure and longer stimulus duration case were verified using voltage clamp electrophysiology (Fig. 3f) and shows an initial spike corresponding to the delivery of the ultrasound stimulus, followed by oscillations.

## 3 Discussion

We model how ultrasound results in membrane deflection and eventually leads to transmembrane voltage changes. In a first, we demonstrate real-time membrane deflection due to ultrasound using high-speed DHM imaging (Supplementary Videos 1 and 2). We leverage the Hodgkin-Huxley equations, which are a set of phenomenological equations describing action potential generation in a squid axon and are one of the most important neuronal models. However, observations of mechanical deflection accompanying action potentials [35] show that the underlying assumptions of the Hodgkin-Huxley model may need to be revisited, as there are mechanical phenomena involved. In the context of ultrasound neuromodulation, our model presents insights into the the generation of action potentials due to mechanical deflections and is theoretically supported by models such as the ones put forth in the past few years [28, 47]. The deflection due to the applied ultrasound stimulus results in a net area change of the membrane between the two pin locations that represent an adherent cell. The area changes take place elastically while maintaining constant volume. This results in a change in capacitance that, when incorporated in the Hodgkin-Huxley model, results in transmembrane voltage changes. Capacitance of the membrane can be modeled using an expression for a parallel plate capacitor [48], and an increase in area results in a proportional increase in capacitance [methods].

The model does not take into account restoring effects of the actin cytoskeleton, whose influence will lower the membrane deflection and cause the inner leaflet to deflect less in than the outer leaflet. This cannot account for the deflection observed experimentally in this work, and only plays a minor role in bringing about capacitance changes as suggested in previous studies [23]. The model and the use of DHM imaging presents opportunities for exploring the influence of ultrasound on native neurons and HEK293 cells, as presented here. It can also be used to image cells that have been engineered to express membrane proteins that are sensitive to ultrasound stimuli, in other words using sonogenetics [49]. At a cellular level, there are two proposed models for the activation of mechanically-gated ion channels: the force from lipid model and the force from filament model. The force from lipid model was put forth by Martinac *et al*. [50] and proposes that changes in membrane tension or local membrane curvature result in opening or closing of channels. In the force from filament model [43], the stimulus is transferred to tethers that connect the membrane to the cytoskeleton. Conformational changes in the tethers result in opening or closing of the channel. In reality, both models play a part in opening and closing a given channel.

Although it is difficult to estimate the relative contribution of these mechanisms, it is possible to estimate the deflection of the cell membrane as highlighted in the preceding sections. This is of particular significance when we consider the membrane-bound proteins such as TRPA1, MsCL [51], Piezo [52] and their interaction with the actin network. Disruption of the actin cytoskeleton has been shown to reduce mechanosensitive activity of such ion channels [53] and it is possibly due to decreased separation between the leaflets of the bilayer when the actin network is disrupted. In addition to quantifying the deflection due to mechanosensitive proteins, there is potential for the system to quantify the forces on the cell due to ultrasound using FRET (Förster resonance energy transfer) force sensors [54].

Our model also predicts the generation of action potentials due to capacitive changes that occur when the adherent cell is exposed to ultrasound. Charge across the membrane is maintained by a gradient in ion concentration across the cell membrane; with Na^+^ ions on the outside and Cl^-^ ions on the inside resulting in a net negative resting potential. As the membrane deflects, it is partially constrained by the adherent regions, resulting in an increase in area of the membrane between the adherent locations. An increase in area results in access to a greater number of cations outside the membrane, driving up the capacitance. This relationship between area, capacitance and transmembrane voltage change has been indicated in prior publications that investigate capacitive properties of biological membranes [48], outside the context of ultrasound neuromodulation. We demonstrate transmembrane voltage changes for two cases, a pressure of 0.5 MPa and 1 MPa and observe that voltage changes only take place for the higher pressure case for lower stimulus durations. In addition to obtaining a pressure threshold at a specific frequency, we also investigate the influence of longer stimulus durations on the generation of action potentials for different cases of baseline capacitance. As verified by a current clamp electrophysiology study in the whole cell configuration, increased stimulus durations even at at lower pressures result in action potential generation, though with lower spike rates.

One of the limitations with performing single cell current clamp electrophysiology is the loss of a seal with the membrane due to the deflection of the membrane in response to ultrasound at amplitudes sufficient to drive a physiological response with respect to the patch pipette. There are, however, reports of current clamp electrophysiology results with ultrasound using microbubbles [55] and at much higher frequencies [56] or with devices [57]. In each of these three cases, there is reason to believe that while the stimulation techniques or device may work for *in vitro* work, they will not be suitable for *in vivo* work. One potential way to overcome this issue would be to perform electrophysiological recordings for cells encased in matrigel that would limit the movement of the recording pipette with respect to the membrane.

Until now, the mechanisms underlying ultrasound neuromodulation have lacked explanation and existing models lack experimental data. Taken together, our results offer valuable insight into the underlying effects of ultrasound on cell membranes, as well as insight into how these effects translate to transmembrane voltage changes. The predictions of our model were confirmed using a novel, high-speed imaging technique. We were able to visualize and quantify membrane deflection in real-time and predict depolarization due to the imposed ultrasound stimulus.

## 4 Methods

### 4.1 HEK293 cell culture

Human embryonic kidney (HEK293) cells (ATCC CRL-1573, Manassas, Virginia, USA) were cultured using standard procedure in DMEM supplemented with 10% fetal bovine serum (FBS) and 20 mM glutamine in a 37°C and 5% CO_2_ incubator. Cells beyond passage 30 were discarded and a new aliquot was thawed. For experimental plating, 18 mm coverslips were coated with poly-d-lysine (PDL; 10 g/L, minimum 2 hours, P6407, Sigma-Aldrich, St. Louis, Missouri, USA), and HEK293 cells were seeded at 150K, 200K, or 250K cells/mL for 24 hours before the experiment. Cells were allowed to grow 24 hours. For imaging, coverslips were mounted on a specialized chamber featuring an ultrasound transducer approximately 2 mm below the coverslip and a 10 mL reservoir of media above the coverslip. Once cells were in focus, a 6.72 MHz ultrasound pulse of 50 ms duration was delivered while imaging with an immersion objective as described in following sections, and a cell membrane profile was reconstructed and analyzed.

### 4.2 Digital holographic microscopy

HEK cells and neurons were observed through a 40X, 0.8 NA (numerical aperture) water immersion microscope objective. The field of view used for the setup was 60.5 *µ*m × 60.5 *µ*m, with a vertical accuracy and repeatability of 4 nm and 0.08 nm respectively [58]. Holograms were recorded using a high-speed camera (Nova S12, Photron, San Diego, California, USA). Acquisition and reconstruction were performed using custom software (Koala, Lynceé-tec Inc., Lausanne, Switzerland) on a computer workstation. Data were recorded on a separate computer equipped with a solid-state drive, with each 100 ms recording equating to∼ 20 Gigabytes of data. The observations reported in this study represent a combined analysis of 1.4 Terabytes of data. The data were reconstructed after each batch of six coverslips was processed in order to reduce the time between trials and to ensure optimum cell health. The setup consisted of a custom perfusion chamber that was built to accommodate a lithium niobate transducer operating at 6.72 MHz. The perfusion chamber was housed on a stage maintained at 37°C (Fig. 1a) using a heated stage (Bioscience Tools TC-100s).

### 4.3 Modeling of deflection and transmembrane voltage changes

As the pressure wave propagates through the fluid and contacts the adherent cell, the region of the cell membrane between adhesion zones deflects. This deflection leads to a change in area of the membrane and causes a capacitance change. The two-dimensional model assumes that the membrane has a known value of surface tension [59]. The membrane is surrounded by a fluid, assumed to have the properties of water in this case. The vertical displacement of the membrane is approximated to be equal to the displacement of the fluid just above the membrane. We start with a simplified version of the Navier-Stokes equation,

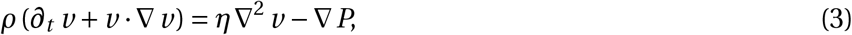

where *ρ* and *η* are the density and viscosity of water, respectively. The expression Δ*P* is the pressure gradient and *v* is the velocity. In Eq. (3), the convective acceleration is *v*· Δ *v =* 0 as the flow is unidirectional in *z* [60] and the fluid is assumed to be incompressible. The membrane is symmetric in *x* and *y*, allowing the viscous term to be simplified as *∂*_*x*_ *v*_*z*_ *= ∂*_*y*_ *v*_*z*_. We are left with

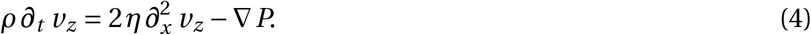

The net pressure gradient in this case is a function of the time dependent pressure in the fluid due to ultrasound and the surface tension of the membrane, which resists deformation:

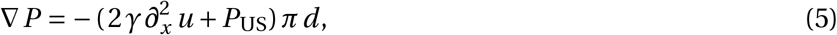

where *u* is the displacement in *z* and *P*_US_ is the pressure due to an ultrasound source, typically acting in the form of a sinusoidal pulse, *P*_US_=*P*_0_ sin(*ωt*), where *ω* =2*πf* . By contrast, Prieto *et al*. [37] at this point chose to represent the ultrasound as a step change in the pressure, from a static, zero relative pressure to a static positive value at time *t* = 0 well below the pressure amplitudes used in experimental studies, typically 1 kPa to 1 MPa. Prieto *et al*.’s representation is numerically attractive but difficult to reconcile with the harmonic oscillatory pressure delivered by the transducer. In the absence of an analytical solution for the ultrasound propagating through the medium and membrane, one would be forced to numerically represent the MHz-order sinusoidal signal with sufficiently small spatiotemporal step sizes to satisfy the Nyquist criterion, and do so for at least several hundred milliseconds to determine the response of the cell membrane to the ultrasound pressure oscillation, representing very large models with many millions to billions of temporal steps for a single solution. Consequently, these past studies have been understandably forced to make spurious approximations^1^ to avoid impossibly prohibitive computation time.

Substituting this into Eq. (4) produces a partial differential equation for the displacement of the membrane driven by ultrasound:

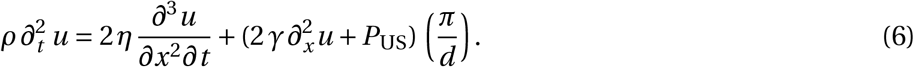

The boundary conditions are the clamped conditions at the ends of the membrane and the initial displacement condition,

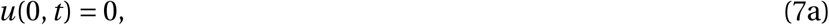

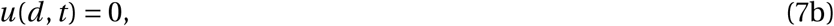

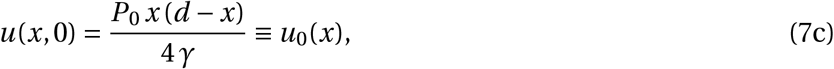

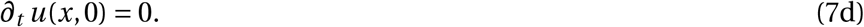

If hydrostatic pressure is included, the initial condition for membrane displacement may be found by solving 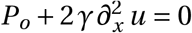. The general soltuion to partial differential Eq. 6 was obtained with the method of eigenfunction expansion, as outlined further on. This is achieved using an orthogonal eigenbasis:

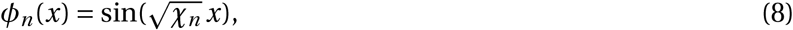

where *χ*_*n*_*=* (*n π*/*d*)^2^ corresponds to the *n*^th^ wavemode for a membrane with diameter *d* . Expanding *u* gives us

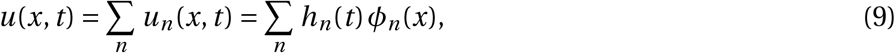

so that clearly the even modes vanish and we may write *n =*2 *k*+ 1, and *k*∈ ℤ≥ 0 where ℤ is an integer set. Substituting this expression into (6), one has

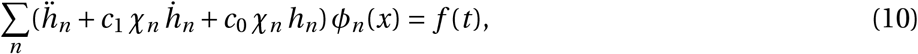

where *c*_1_= 2 *η*/*ρ* and *c*_0_ = 2 *πγ*/*ρ d*, are written in terms of the density of the surrounding fluid, *ρ*; the viscosity of the surrounding fluid, *η*; the surface tension along the fluid-membrane interface, *γ*; and the membrane diameter, *d* . By multiplying both sides by φ _*m*_(*x*) (with *m∈* ℤ^+^), integrating over *x* from 0 to *d*, and then leveraging the orthogonality of sines, we find that the time-dependent component for the *n*^th^ eigenmode satisfies the second-order ordinary differential equation

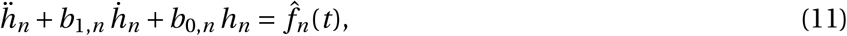

where *b*_1,*n*_ *= c*_1_ *χ*_*n*_, *b*_0,*n*_ *= c*_0_ *χ*_*n*_, and

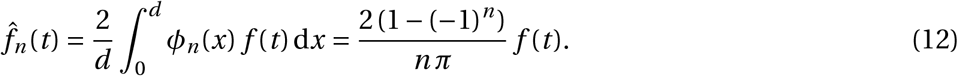

The means for obtaining a solution to equations of the form (11) is well known. The homogeneous solution and its coefficients are given by

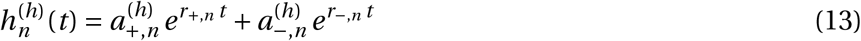

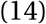

where the coefficients 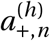 and 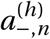 are

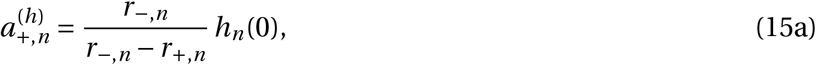

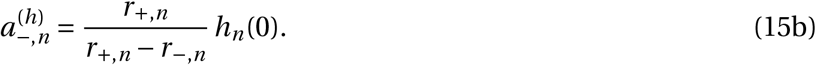

The inhomogeneous solution is

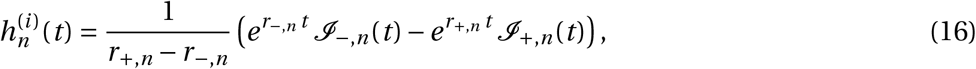

where

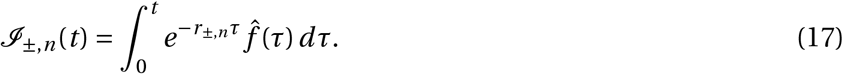

The total waveform solution is then numerically implemented by taking a finite-term approximation of (9). The change in area, *A*, of the membrane then be calculated once the time-dependent membrane deflection is obtained:

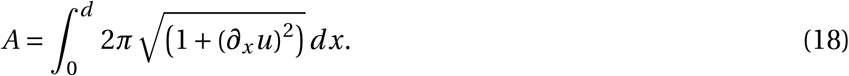

By extension, this allows us to determine the change in membrane capacitance, *C*, due to the area change,

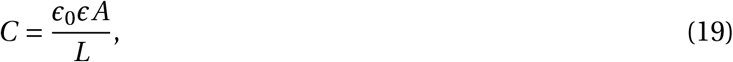

where we have regarded the membrane as a dielectric between two charged surfaces. In this case, *L* is the thickness of the bilayer and has values between 4 nm and 9 nm, and the relative permittivity, *∈*, has a value of 2 [61].

The above value of capacitance change is coupled with the modified Hodgkin-Huxley neuronal model, where the capacitive current is defined as 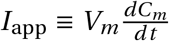. This model contains a voltage-gated sodium current and delayed-rectifier potassium current to generate actions, a slow non-inactivating potassium current to recapitulate the spike-frequency adaptation behavior seen in thalamocortical cells, and a leakage current.

Equation (20) defines the voltage-gated Na^+^ current where 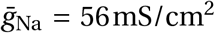 is the maximal conductance and *E*_Na_ = 50 mV is the Nernst potential of the Na^+^ channels. The parameter *V*_th_= − 56.2 mV sets the spike threshold

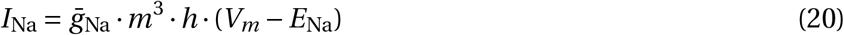

where the gating variables *m* and *h* vary with time according to

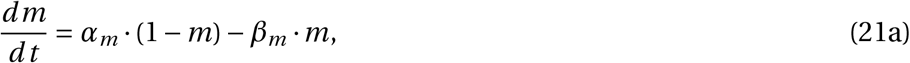

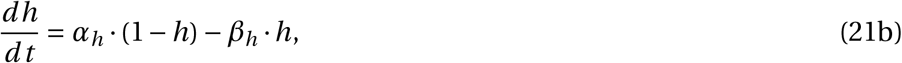

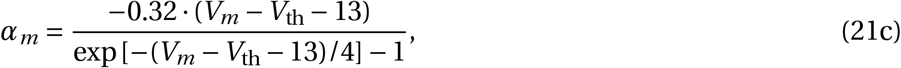

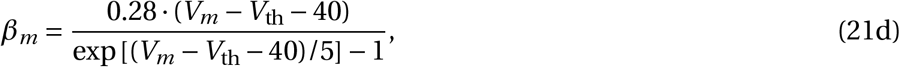

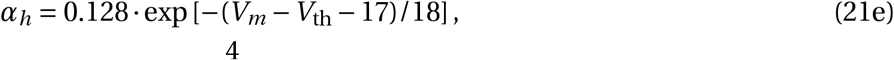

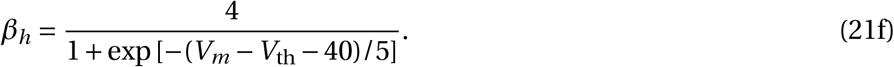

The delayed rectifier K^+^ current is

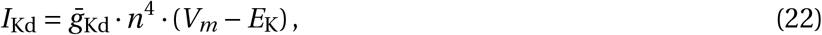

where 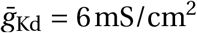 is the maximal conductance of the delayed-rectifier K^+^ channels and *E*_K_ = − 90 mV is the Nernst potential of the K^+^ channels, and with *n* evolving over time as

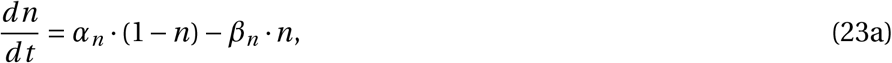

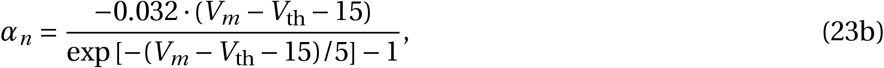

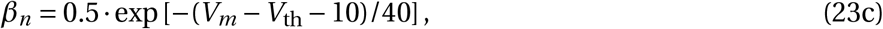

A slow non-inactivating K^+^ current may be defined as

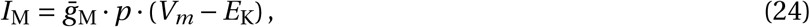

where 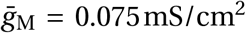 is the maximal conductance and *τ*_max_ = 608 ms is the decay time constant for adaptation of the slow non-inactivation K^+^ channels. The parameter *p* is such that

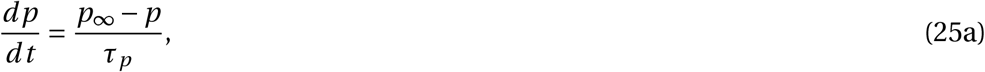

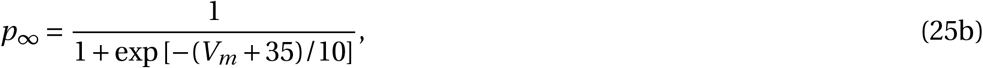

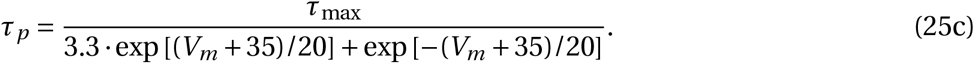

The leakage current is

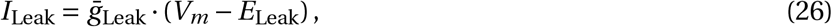

where 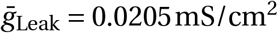 is the maximal conductance and *E*_Leak_= − 70.3 mV is the Nernst potential of the non-voltage-dependent, non-specific ion channels.

We set the following initial conditions for the gating terms:

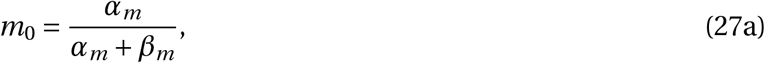

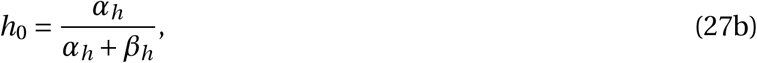

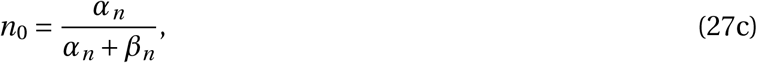

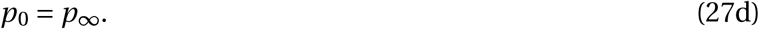

Equations (20) through (25) are solved with initial conditions (27) to obtain the transmembrane voltage change of a neuron when subjected to ultrasound stimuli.

### 4.4 Sustaining oscillations on the membrane

A better understanding of the membrane wave propagation can be obtained by considering the decay transience of the constituent wavemodes within the context of the solution to Eq. (11). Each wavemode will have a solution of the form

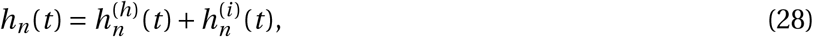

where 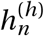 is the homogeneous solution and 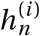 is the inhomogeneous solution for the forced wavemode propagation initialized from zero initial conditions. The general form of the former can be used to characterize the decay transience,

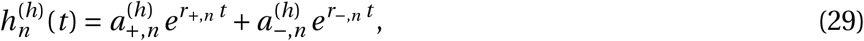

where the coefficients 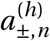 are determined by the initial conditions and *r*_*±,n*_ are the eigenvalues of the left side of (11) (the roots of the characteristic equation):

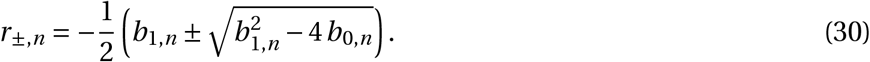

Then the discriminant determines the character of the wavemode:

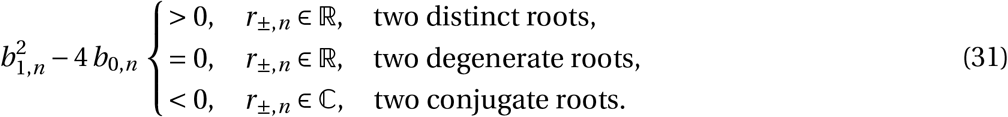

The physical conditions for degeneracy require an exacting degree of marginality rarely (if ever) encountered in real systems, so that we may safely ignore this solution type (degeneracy corresponds to algebraic growth at small times that is mediated by exponential decay at long times).

Rewriting the conditions (31) in terms of physical parameters, one finds that

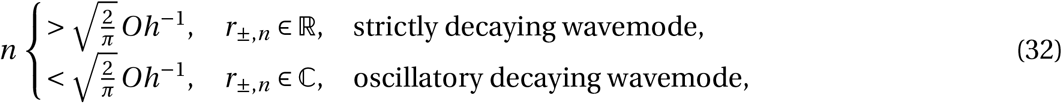

where

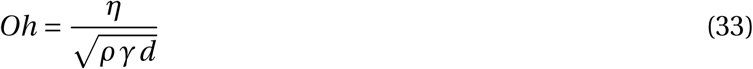

is the Ohnesorge number characterizing the balance between the dissipative viscous effects and the conservative effects resulting from interaction between inertia and surface tension. There exists a condition for oscillation of the unforced membrane and this condition is 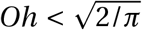. When 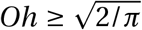, no oscillatory unforced wavemodes are permitted and the unforced membrane *will not oscillate*. When the condition *is* satisfied, one observes that oscillation can be attributed exclusively to wavemodes with the “smallest” mode numbers, and that these will always include the fundamental mode. Fig. S3 represents the change in *Oh* for a range of surface tensions and membrane length.

### 4.5 Ultrasound transducer fabrication

We used a set of custom-made single crystalline 127.86 Y-rotated X-propagating lithium niobate transducers operating in the thickness mode, as described previously [62]. The fundamental frequency was measured to be 6.72 MHz using non-contact laser Doppler vibrometry (UHF-120SV, Polytec, Waldbronn, Germany). The transducers were coated with a 1 *µ*m layer of Au atop 20 nm of Ti acting as an adhesion layer, using a direct-current sputtering (Denton 635 DC Sputtering system) process was used to coat 4 inch diameter wafers in an inert gas environment with a 2.3 mTorr pressure and rotation speed of 13 rpm, at a deposition rate of 1.5 A/s for Ti and 7 A/s for Au. Devices were diced to size (12 mm× 12 mm) and built in to the *in vitro* test setup using an automated dicing saw (DISCO 3220, DISCO, Tokyo Japan).

### 4.6 Rat Primary Neuron Culture

Rat primary neuronal cultures were prepared from rat pup tissue at embryonic days (E) 18 containing combined cortex, hippocampus and ventricular zone. The tissue was obtained from BrainBits (Catalog #: SDE-HCV) in Hibernate-E media and used the same day for dissociation following their protocol.

Briefly, tissue was incubated in a solution of papain (BrainBits PAP) at 2 mg/mL for 30 min at 37°C and dissociated in Hibernate-E for one minute using one sterile 9” silanized Pasteur pipette with a fire-polished tip. The cell dispersion solution was centrifuged at 1100 rpm for 1 min, and the pellet was resuspended with 1 mL NbActiv1 (BrainBits NbActiv1 500 mL). The cell concentration was determined using a haemocytometer (TC20, Bio-Rad Labs, Hercules, California, USA) and neurons were plated in 12-well culture plates with 18-mm PDL-coated coverslips (GG-18-PDL, Neuvitro Corporation, Vancouver, Washington, USA) at a concentration of 1.3 million cells/well. Neurons were then incubated at 37°C, 5% CO_2_, performing half media changes every 3-4 days with fresh NbActiv1 supplemented with PrimocinTM (ant-pm-1, InvivoGen, San Diego, California, USA). Cultures were incubated at 37°C, 5% CO_2_ until day 10–12 and were used in DHM imaging experiments.

### 4.7 *In-vitro* electrophysiology

A stable line of neurons using the protocol listed above were cultured on 18 mm round coverslips, at a seeding density of ∼ 300k cells/well in a tissue-culture treated 12-well plate. Neurons were allowed to mature for 11-14 days in vitro prior to recording. Coverslips were transferred to a custom machined acrylic stage containing a bath of external solution; NaCl (140 mM), KCl (4 mM), MgCl_2_ (2 mM), glucose (5 mM), and HEPES (10 mM) with an osmolarity of ∼ 290 mOsm. Patch pipettes were pulled on a pipette puller (P-97, Sutter Instruments, Novato, CA, USA) programmed to give 4-6 MΩ tips from filamented borosilicate glass (o.d. 1.5 mm, i.d. 0.86 mm) and used with an internal solution comprising of a CsF and KF base (#08 3008, #08 3007, Nanion, Munich, Germany). A 40X water dipping lens (LUMPLFLN40XW, Olympus Corporation, Tokyo, Japan) with 0.8 NA was used in combination with a cMOS camera (01-OPTIMOS-R-M-16-C QImaging OptiMOS, Roper Technologies, USA) to visualize cells with Köhler or fluorescent illumination. Electrical signals were acquired using an amplifier (Axon Instruments Multiclamp 700B, Molecular Devices LLC, California, USA) and digitized (Axon Instruments Digidata 1550B, Molecular Devices LLC, California, USA) using an acquisition and control software (pClamp 11, Molecular Devices LLC, California, USA). Gap free recordings were conducted (typically holding the membrane potential at− 70 mV) while delivering the ultrasound stimulus. The ultrasound delivery rig used for patch clamp experiments was the same used for imaging experiments. Briefly, waveforms were programmed using an arbitrary function generator (33600A Series, Keysight, California, USA) connected via BNC to an amplifier (TC2057574, Vox Technologies, Richardson, TX). Military communications grade BNC cables (CA5512-36, Federal Custom Cable, California, USA) were used to ensure impedance matching in our systems and reduce electrical interference. The amplifier was connected to our custom-made lithium niobate transducer mounted on a dovetail sliding arm, and coupled to the bottom of the recording chamber with ultrasound gel. Recordings were carried out in response to peak pressures of 0.5 MPa as access resistance could not be maintained when high pressures were delivered. Upon successful whole-cell access, baseline gap-free recordings in current clamp trials were obtained. Access resistance during successful whole-cell recordings was maintained between 10 to 25 MΩ.

## Acknowledgements

J.F. is grateful for funding for this work from the W.M. Keck Foundation via a SERF grant and, with S.C., from the National Institutes of Health (NIH) via R01NS115591. S.C. is also grateful to the NIH in support of this work via grant R01MH MH111534. J.O. is thankful for support provided by the University of California’s Presidential Postdoctoral Fellowship Program. This work was performed at the Medically Advanced Devices Laboratory at the University of California, San Diego. Fabrication was performed in part at the San Diego Nanotechnology Infrastructure (SDNI) of UCSD, a member of the National Nanotechnology Coordinated Infrastructure, which is supported by the National Science Foundation (Grant ECCS–1542148). The authors would like to thank members of the Medically Advanced Devices Laboratory for feedback, and Prof. Juan Lasheras^†^ and Dr. Yi-Ting Yeh for use of their cell culture equipment. The authors would also like to thank Benjamin Rappaz, Frank Liu and Yves Emery from Lynceé-tec for designing and configuring the bespoke high-speed digital holographic microscope for this work.

Most importantly, the authors wish to dedicate this paper to the memory of Prof. Juan Lasheras.

## Conflict of Interest

The authors declare no conflict of interest.

## Author contributions

A.V., S.C. and J.F. designed the experiments. A.V. conducted the experiments. A.V. developed the model with support from J.O. and J.F. U.M. and C.W. performed the HEK cell culture. M.D. and A.V. performed the primary neuron culture. Y.T. performed the electrophysiology. A.V. wrote the paper with edits from J.F., J.O., and S.C.

## Footnotes

1 One can use the analytical solution to show that the results from the noted study are spurious when the correct amplitudes and frequencies of the ultrasound are used.

## Notes

### Competing Interest Statement

The authors have declared no competing interest.

## References

[1] H. S. Mayberg, A. M. Lozano, V. Voon, H. E. McNeely, D. Seminowicz, C. Hamani, J. M. Schwalb, S. H. Kennedy, Neuron 2005, 45, 5 651.

[2] J. M. Bronstein, M. Tagliati, R. L. Alterman, A. M. Lozano, J. Volkmann, A. Stefani, F. B. Horak, M. S. Okun, K. D. Foote, P. Krack, et al., Archives of Neurology 2011, 68, 2 165.

[3] D. A. Groves, V. J. Brown, Neuroscience & Biobehavioral Reviews 2005, 29, 3 493.

[4] J. De Kroon, J. Van der Lee, M. J. IJzerman, G. Lankhorst, Clinical Rehabilitation 2002, 16, 4 350.

[5] M. Ranjan, A. Boutet, S. Bhatia, A. Wilfong, W. Hader, M. R. Lee, A. R. Rezai, P. D. Adelson, Expert Review of Neurotherapeutics 2019, 19, 10 937.

[6] B. Y. Chow, E. S. Boyden, Science Translational Medicine 2013, 5, 177 177ps5.

[7] J. G. English, B. L. Roth, JAMA Neurology 2015, 72, 11 1361.

[8] V. Busskamp, S. Picaud, J.-A. Sahel, B. Roska, Gene Therapy 2012, 19, 2 169.

[9] L. Fenno, O. Yizhar, K. Deisseroth, Annual Review of Neuroscience 2011, 34.

[10] H. H. Yoon, J. H. Park, Y. H. Kim, J. Min, E. Hwang, C. J. Lee, J.-K. Francis Suh, O. Hwang, S. R. Jeon, Neurosurgery 2014, 74, 5 533.

[11] W. M. Grill, S. E. Norman, R. V. Bellamkonda, Annual Review of Biomedical Engineering 2009, 11 1.

[12] J. W. Salatino, K. A. Ludwig, T. D. Y. Kozai, E. K. Purcell, Nature Biomedical Engineering 2017, 1, 11 862.

[13] A. S. Tolias, F. Sultan, M. Augath, A. Oeltermann, E. J. Tehovnik, P. H. Schiller, N. K. Logothetis, Neuron 2005, 48, 6 901.

[14] T. Wagner, A. Valero-Cabre, A. Pascual-Leone, Annu. Rev. Biomed. Eng. 2007, 9 527.

[15] A. T. Barker, Electroencephalogr Clin Neurophysiol Suppl. 1999, 51 3.

[16] E. S. Boyden, F. Zhang, E. Bamberg, G. Nagel, K. Deisseroth, Nature Neuroscience 2005, 8, 9 1263.

[17] P. Aldrin-Kirk, T. Björklund, In Viral Vectors for Gene Therapy, 59–87. Springer, 2019.

[18] M. Duque, C. A. Lee-Kubli, Y. Tufail, U. Magaram, J. M. Lopez, E. Edsinger, A. Vasan, R. Shiao, C. Weiss, J. Friend, S. H. Chalasani, bioRxiv 2020.

[19] J. O. Szablowski, A. Bar-Zion, M. G. Shapiro, Accounts of Chemical Research 2019, 52, 9 2427.

[20] J. Blackmore, S. Shrivastava, J. Sallet, C. R. Butler, R. O. Cleveland, Ultrasound in Medicine & Biology 2019, 45, 7 1509.

[21] E. N. Harvey, American Journal of Physiology-Legacy Content 1929, 91, 1 284.

[22] T. Yang, J. Chen, B. Yan, D. Zhou, Medical Hypotheses 2011, 76, 3 381.

[23] M. Plaksin, S. Shoham, E. Kimmel, Physical Review X 2014, 4, 1 011004.

[24] H. Guo, M. Hamilton II, S. J. Offutt, C. D. Gloeckner, T. Li, Y. Kim, W. Legon, J. K. Alford, H. H. Lim, Neuron 2018, 98, 5 1020.

[25] F. Di Giacinto, M. De Spirito, G. Maulucci, Ultrasound in Medicine & Biology 2019, 45, 5 1143.

[26] K. Iwasa, I. Tasaki, R. C. Gibbons, Science 1980, 210, 4467 338.

[27] Z. Gil, S. D. Silberberg, K. L. Magleby, Proceedings of the National Academy of Sciences 1999, 96, 25 14594.

[28] T. Heimburg, A. D. Jackson, Proceedings of the National Academy of Sciences 2005, 102, 28 9790.

[29] F. Zernike, Physica 1942, 9, 10 974.

[30] R. Allen, G. David, Zeitschrift für Wissenschaftliche Mikroskopie und Mikroskopische Technik 1969, 69, 4 193.

[31] B. Rappaz, P. Marquet, E. Cuche, Y. Emery, C. Depeursinge, P. J. Magistretti, Optics Express 2005, 13, 23 9361.

[32] U. Schnars, W.P. Jüptner, Measurement Science and Technology 2002, 13, 9 R85.

[33] F. Dubois, L. Joannes, J.-C. Legros, Applied Optics 1999, 38, 34 7085.

[34] A. Stadelmaier, J. H. Massig, Optics Letters 2000, 25, 22 1630.

[35] G. Kim, P. Kosterin, A. Obaid, B. Salzberg, Biophysical Journal 2007, 92, 9 3122.

[36] P. L. Frederix, P. D. Bosshart, A. Engel, Biophysical Journal 2009, 96, 2 329.

[37] M. L. Prieto, Ö. Oralkan, B. T. Khuri-Yakub, M. C. Maduke, PLoS One 2013, 8, 10 e77115.

[38] M. Israelowitz, B. Weyand, S. Rizvi, P. M. Vogt, H. P. von Schroeder, Journal of Healthcare Engineering 2012, 3, 3 455.

[39] Y. Guyot, F. Luyten, J. Schrooten, I. Papantoniou, L. Geris, Biotechnology and Bioengineering 2015, 112, 12 2591.

[40] C. K. Holland, R. E. Apfel, IEEE Transactions on Ultrasonics, Ferroelectrics, and Frequency Control 1989, 36, 2 204.

[41] C. C. Church, C. Labuda, K. Nightingale, Ultrasound in Medicine and Biology 2015, 41, 2 472.

[42] American Institute of Ultrasound in Medicine, Statement on mammalian biological effects in tissues without gas bodies, 2015.

[43] C. D. Cox, N. Bavi, B. Martinac, In Current Topics in Membranes, volume 79, 59–96. Elsevier, 2017.

[44] A. L. Hodgkin, A. F. Huxley, The Journal of Physiology 1952, 117, 4 500.

[45] L. Abbott, T. B. Kepler, In Statistical mechanics of neural networks, 5–18. Springer, 1990.

[46] G. Eyal, M. B. Verhoog, G. Testa-Silva, Y. Deitcher, J. C. Lodder, R. Benavides-Piccione, J. Morales, J. DeFelipe, C. P. de Kock, H. D. Mansvelder, et al., Elife 2016, 5 e16553.

[47] H. Chen, D. Garcia-Gonzalez, A. Jérusalem, Physical Review E 2019, 99, 3 032406.

[48] L. D. Mosgaard, K. A. Zecchi, T. Heimburg, Soft Matter 2015, 11, 40 7899.

[49] S. Ibsen, A. Tong, C. Schutt, S. Esener, S. H. Chalasani, Nature communications 2015, 6, 1 1.

[50] B. Martinac, J. Adler, C. Kung, Nature 1990, 348, 6298 261.

[51] J. Ye, S. Tang, L. Meng, X. Li, X. Wen, S. Chen, L. Niu, X. Li, W. Qiu, H. Hu, et al., Nano letters 2018, 18, 7 4148.

[52] Z. Qiu, J. Guo, S. Kala, J. Zhu, Q. Xian, W. Qiu, G. Li, T. Zhu, L. Meng, R. Zhang, et al., iScience 2019, 21 448.

[53] K. Hayakawa, H. Tatsumi, M. Sokabe, Journal of Cell Science 2008, 121, 4 496.

[54] J. Guo, F. Sachs, F. Meng, Antioxidants & Redox Signaling 2014, 20, 6 986.

[55] Z. Fan, H. Liu, M. Mayer, C. X. Deng, Proceedings of the National Academy of Sciences of the United States of America 2012, 109, 41 16486.

[56] P. S. Balasubramanian, A. Singh, C. Xu, A. Lal, Scientific Reports 2020, 10, 1 3075.

[57] Z. Lin, W. Zhou, X. Huang, K. Wang, J. Tang, L. Niu, L. Meng, H. Zheng, Advanced Biosystems 2018, 2, 8 1800041.

[58] Lyncéetec, Specifications of high-speed digital holographic imaging system, https://www.lynceetec.com/high-speed-dhm/#tab-3, 2020.

[59] F. Jähnig, Biophysical Journal 1996, 71, 3 1348.

[60] N. Riley, Annual Review of Fluid Mechanics 2001, 33, 1 43.

[61] W.-T. Huang, D. Levitt, Biophysical Journal 1977, 17, 2 111.

[62] A. Vasan, W. Connacher, J. Friend, Journal of Visualized Experiments 2020.

